# A deep neural network approach to predicting clinical outcomes of neuroblastoma patients

**DOI:** 10.1101/750364

**Authors:** Léon-Charles Tranchevent, Francisco Azuaje, Jagath C. Rajapakse

## Abstract

The availability of high-throughput omics datasets from large patient cohorts has allowed the development of methods that aim at predicting patient clinical outcomes, such as survival and disease recurrence. Such methods are also important to better understand the biological mechanisms underlying disease etiology and development, as well as treatment responses. Recently, different predictive models, relying on distinct algorithms (including Support Vector Machines and Random Forests) have been investigated. In this context, deep learning strategies are of special interest due to their demonstrated superior performance over a wide range of problems and datasets. One of the main challenges of such strategies is the “small n large p” problem. Indeed, omics datasets typically consist of small numbers of samples and large numbers of features relative to typical deep learning datasets. Neural networks usually tackle this problem through feature selection or by including additional constraints during the learning process.

We propose to tackle this problem with a novel strategy that relies on a graph-based method for feature extraction, coupled with a deep neural network for clinical outcome prediction. The omics data are first represented as graphs whose nodes represent patients, and edges represent correlations between the patients’ omics profiles. Topological features, such as centralities, are then extracted from these graphs for every node. Lastly, these features are used as input to train and test various classifiers.

We apply this strategy to four neuroblastoma datasets and observe that models based on neural networks are more accurate than state of the art models (DNN: 85%-87%, SVM/RF: 75%-82%). We explore how different parameters and configurations are selected in order to overcome the effects of the small data problem as well as the curse of dimensionality. Our results indicate that the deep neural networks capture complex features in the data that help predicting patient clinical outcomes.

## Background

A lot of efforts have been made recently to create and validate predictive models for clinical research. In particular, the identification of relevant biomarkers for diagnosis and prognosis has been facilitated by the generation of large scale omics datasets for large patient cohorts. Candidate biomarkers are now identified by looking at all bioentities, including non-coding transcripts such as miRNA [31, 13], in different tissues, including blood [23, 21] and by investigating different possible levels of regulation, for instance epigenetics [19, 10, 29].

One challenging objective is to identify prognostic biomarkers, *i.e.*, biomarkers that can be used to predict the clinical outcome of patients such as whether the disease will progress or whether the patient will respond to a treatment. One strategy to identify such biomarkers is to build classifiers that can effectively classify patients into clinically relevant categories. For instance, various machine learning models predicting the progression of the disease and even the death of patients were proposed for neuroblastoma [37]. Similar models have also been built for other diseases such as ovarian cancer to predict the patients’ response to chemotherapy using different variants of classical learning algorithms such as Support Vector Machines (SVM) and Random Forest (RF) [34]. More recently, gynecologic and breast cancers were classified into five clinically relevant subtypes based on the patients extensive omics profiles extracted from The Cancer Genome Atlas (TCGA)[3]. A simple decision tree was then proposed to classify samples and thus predict the clinical outcome of the associated patients. Although the general performance of these models is encouraging, they still need to be improved before being effectively useful in practice.

This study aims at improving these approaches by investigating a graph-based feature extraction method, coupled with a deep neural network, for patient clinical outcome prediction. One challenge when applying a machine learning strategy to omics data resides in the properties of the input data. Canonical datasets usually contain many instances but relatively few attributes. In contrast, biomedical datasets such as patient omics datasets usually have a relatively low number of instances (*i.e.*, few samples) and a relatively high number of attributes (*i.e.*, curse of dimensionality). For instance, the large data repository TCGA contains data for more than 11,000 cancer patients, and although the numbers vary from one cancer to another, for each patient, a least a few dozens of thousands of attributes are available [24]. The situation is even worse when focusing on a single disease or phenotype, for which less than 1,000 patients might have been screened [4, 6, 11].

Previous approaches to handle omics data (with few samples and many features) rely on either feature selection via dimension reduction [18, 12, 30] or on imposing constraints on the learning algorithm [17, 9]. For instance, several studies have coupled neural networks to Cox models for survival analysis [33, 14]. These methods either perform feature selection before inputing the data to deep neural network [33, 14] or let the Cox model perform the selection afterwards [32]. More recently, the GEDFN method was introduced, which relies on a deep neural network to perform disease outcome classification [17]. GEDFN handles the curse of dimensionality by imposing a constraint on the first hidden layer. More precisely, a feature graph (in this case, a protein-protein interaction network) is used to enforce sparsity of the connections between the input layer and the first hidden layer.

We propose a strategy to create machine-learning models starting from patient omics datasets by first reducing the number of features through a graph topological analysis. Predictive models can then be trained and tested, and their parameters can be fine-tuned. Due to their high performance on many complex problems involving high-dimensional datasets, we build our approach around Deep Neural Networks (DNN). Our hypothesis is that the complex features explored by these networks can improve the prediction of patient clinical outcomes. We apply this strategy to four neuroblastoma datasets, in which the gene expression levels of hundreds of patients have been measured using different technologies (*i.e.*, microarray and RNA-sequencing). In this context, we investigate the suitability of our approach by comparing it to state of the art methods such as SVM and RF.

## Methods

### Data collection

The neuroblastoma transcriptomics datasets are summarized in Table 1. Briefly, the data were downloaded from GEO [1] using the identifiers GSE49710 (tag ‘*Fischer-M*’), GSE62564 (tag ‘*Fischer-R*’) and GSE3960 (tag ‘*Maris*’). The pre-processed transcriptomics data are extracted from the GEO matrix files for 498 patients (‘*Fischer-M*’ and ‘*Fischer-R*’) and 102 patients (‘*Maris*’). In addition, clinical descriptors are also extracted from the headers of the GEO matrix files (‘*Fischer-M*’ and ‘*Fischer-R*’) or from the associated publications (‘*Maris*’). For ‘*Maris*’, survival data for ten patients are missing, leaving 92 patients for analysis. A fourth dataset (tag ‘*Versteeg*’) is described in GEO record GSE16476. However the associated clinical descriptors are only available through the R2 tool [2]. For consistency, we have also extracted the expression profiles for the 88 patients using the R2 tool. In all four cases, the clinical outcomes include ‘*Death from disease*’ and ‘*Disease progression*’, as binary features (absence or presence of event) which are used to define classes. Genes or transcripts with any missing value are dropped. The number of features remaining after pre-processing are 43,291, 43,827, 12,625 and 40,918 respectively for the ‘*Fischer-M*’, ‘*Fischer-R*’, ‘*Maris*’ and ‘*Versteeg*’ matrices.

**Table 1:** Details about the four expression datasets used in the present study.

| Name | Reference | Data type | Size | Usage |
| --- | --- | --- | --- | --- |
| *Fischer-M* | Zhang *et al.*, 2014 [37, 27] | Microarray | 498 * 43,291 | Training, testing |
| *Fischer-R* | Zhang *et al.*, 2014 [37, 27] | RNA-seq | 498 * 43,827 | Training, testing |
| *Maris* | Wang *et al.*, 2006 [28] | Microarray | 92 * 12,625 | Testing |
| *Versteeg* | Molenar *et al.*, 2012 [20] | Microarray | 88 * 40,918 | Testing |

### Data processing through topological analysis

Each dataset is then reduced through a Wilcoxon analysis that identifies the features (*i.e.*, genes or transcripts) that are most correlated with each clinical outcome using only the training data (Wilcoxon *P <* 0.05). When this analysis did not return any feature, the top 5% features were used regardless of their p-values (for *Maris*’ and ‘*Versteeg*’). After dimension reduction, there are between 638 and 2,196 features left depending on the dataset and the clinical outcome.

These reduced datasets are then used to infer Patient Similarity Networks (PSN), graphs in which a node represents a patient and an edge between two nodes represents the similarity between the two profiles of the corresponding patients. These graphs are built first, by computing the Pearson correlation coefficients between all profiles pairwise and second, by normalizing and rescaling these coefficients into positive edge weights through a WGCNA analysis [36], as described previously [25]. These graphs contain one node per patient, are fully connected and their weighted degree distributions follow a power law (*i.e.*, scale-free graphs). Only one graph is derived per dataset, and each of the four datasets is analyzed independently. This means that for ‘*Fischer*’ datasets, the graph contains both training and testing samples.

Various topological features are then extracted from the graphs, and will be used to build classifiers. In particular, we compute twelve centrality metrics as described previously (weighted degree, closeness centrality, current-flow closeness centrality, current-flow betweenness centrality, eigen vector centrality, Katz centrality, hit centrality, page-rank centrality, load centrality, local clustering coefficient, iterative weighted degree and iterative local clustering coefficient) for all four datasets. In addition, we perform clustering of each graph using spectral clustering [26] and Stochastic Block Models (SBM) [7]. The optimal number of modules is determined automatically using dedicated methods from the spectral clustering and SBM packages. For the two ‘*Fischer*’ datasets and the two clinical outcomes, the optimal number of modules varies between 5 and 10 for spectral clustering and 25 and 42 for SBM. This analysis was not performed for the other datasets. All repartitions are used to create modularity features. Each modularity feature represents one single module and is binary (its value is set to one for members of the module and zero otherwise). All features are normalized before being feed to the classifiers (to have a zero mean and unit variance). Two datasets can be concatenated prior to the model training, all configurations used in this study are summarized in Table 2.

**Table 2:** List of the possible data configurations (topological feature sets, datasets) used to train classification models. ^*a*^ The number of modules for each graph, corresponding to one clinical outcomes of interest, is different. ^*b*^ This is the combined dataset in which the topological features of both ‘*Fischer-M*’ and ‘*Fischer-R*’ are concatenated.

| Datasets | Topological features | Total size |
| --- | --- | --- |
| *Fischer-M* | Centralities | 12 |
|  | Modularities | {30, 39}<sup>a</sup> |
|  | Both | {42, 51}<sup>a</sup> |
| *Fischer-R* | Centralities | 12 |
|  | Modularities | {36, 47}<sup>a</sup> |
|  | Both | {48, 59}<sup>a</sup> |
| *Fischer<sup>b</sup>* | Centralities | 24 |
|  | Modularities | {75, 77}<sup>a</sup> |
|  | Both | {99, 101}<sup>a</sup> |

### Modeling through deep neural networks

Classes are defined by the binary clinical outcomes ‘*Death from disease*’ and ‘*Disease progression*’. For the ‘*Fischer*’ datasets, the original patient stratification [37] is extended to create three groups of samples through stratified sampling: a training set (249 samples, 50%), an evaluation set (125 samples, 25%) and a validation set (124 samples, 25%). The proportions of samples associated to each clinical outcome of interest remain stable among the three groups (Additional File 2).

Deep Neural Networks (DNN) are feed forward neural networks with hidden layers, which can be trained to solve classification and regression problems. The parameters of these networks are represented by the weights connecting neurons and learned using gradient decent techniques. The DNN models are based on a classical architecture with a varying number of fully connected hidden layers of varying sizes. The activation function of all neurons is the rectified linear unit (ReLU). The softmax function is used as the activation function of the output layer. The training is performed by minimizing the cross-entropy loss function. A mini-batches size of 32 samples is used for training (total size of the training set is 249) and models are ran for 1,000 epochs with an evaluation taking place every 10 epochs. Sample weights are introduced to circumvent the unbalance between the classes (the weights are inversely proportional to the class frequencies). To facilitate replications, random seeds are generated and provided to each DNN model. For our application, DNN classifiers with various architectures are trained. First, the number of hidden layers varies between one and four, and the number of neurons per hidden layer also varies from 2 to 8 (∈{2, 4, 8}). Second, additional parameters such as dropout, optimizer and learning rate are also optimized using a grid search. In particular, dropout is set between 15% and 40% (step set to 5%), learning rate between 1*e*-4 and 5*e*-2 and the optimizer is one among adam, adadelta, adagrad and proximal adagrad. Each DNN model is run ten times with different initialization weights and biases.

### Other modeling approaches

For comparison purposes, SVM and RF models are also trained on the same data. The cost (linear SVM), gamma (linear and RBF SVM) and number of trees (RF) parameters are optimized using a grid search. The cost and gamma parameters are set to 2^2*p*^, with 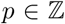, *p* ∈ [−4, 4]. The number of trees varies between 100 and 10,000. Since RF training is non deterministic, the algorithm is run ten times. The SVM optimization problem is however convex and SVM is therefore run only once.

GEDFN accepts omics data as input together with a feature graph. Similarly to the original paper, we use the HINT database v4 [5] to retrieve the human protein-protein interaction network (PPIN) to be used as a feature graph [17]. The mapping between identifiers is performed through BioMart at EnsEMBL v92 [35]. First, the original microarray features (*e.g.*, microarray probesets) are mapped to RefSeq or EnsEMBL transcripts identifiers. The RNA-seq features are already associated to RefSeq transcripts. In the end, transcript identifiers are mapped to UniProt/TrEMBL identifiers (which are the ones also used in the PPIN). The full datasets are too large for GEDFN so the reduced datasets (after dimension reduction) described above are used as inputs. For comparison purposes, only the ‘*Fischer-M*’ data is used for ‘*Death from disease*’ and both ‘*Fischer*’ datasets are concatenated for ‘*Disease progression*’. GEDFN parameter space is explored using a small grid search that always include the default values suggested by the authors. The parameters we optimize are the number of neurons for the second and third layers (∈ {(64, 16), (16, 4)}), the learning rate (∈ {1*e*-4, 1*e*-2}), the adam optimizer regularization (∈{*True, False*}), the number of epochs (∈{100, 1000}) and the batch size (∈{8, 32}). Each GEDFN model is run ten times with different initialization weights and biases. Optimal models for the two clinical outcomes are obtained by training for 1,000 epochs and enforcing regularization.

### Model performance

The performance of each classification model is measured using balanced accuracy (bACC) since the dataset is not balanced (*e.g.*, 4:1 for ‘*Death from disease*’ and 2:1 for ‘*Disease progression*’ in the ‘*Fischer*’ datasets, Additional File 2). In addition, one way ANOVA tests followed by post-hoc Tukey tests are employed for statistical comparisons. We consider p-values smaller than 0.01 as significant. When comparing two conditions, we also consider the difference in their average performance, and the confidence intervals for that difference (noted ∆_*bACC*_). Within any category, the model associated with the best balanced accuracy is considered optimal (including across replicates).

### Implementation

The data processing was performed in python (using packages numpy and pandas). The graph inference and topological analyses were performed in python and C++ (using packages networkx, scipy, igraph, graph-tool and SNFtool). The SVM and RF classifiers were built in R (with packages randomForest and e1071). The DNN classifiers were built in python (with TensorFlow) using the *DNNClassifier* estimator. Training was performed using only CPU cores. GEDFN was run in Python using the implementation provided by the authors. Figures and statistical tests were prepared in R.

## Results

We propose a strategy to build patient classification models, starting from a limited set of patient samples associated with large feature vectors. Our approach relies on a graph-based method to perform dimension reduction by extracting features that are then used for classification (Figure 1 and Methods). Briefly, first the original data are transformed into patient graphs and topological features are extracted from these graphs. These topological features are then used to train deep neural networks. Their classification performance is then compared with those of other classifiers, including Support Vector Machines and Random Forests. We apply this strategy to a previously published cohort of neuroblastoma patients that consist of transcriptomics profiles for 498 patients (‘*Fischer*’, Table 1) [37]. Predictive models are built with a subset of these data and are then used to predict the clinical outcome of patients whose profiles have not been used for training. We then optimize the models and compare their performance by considering their balanced accuracy. The optimal models obtained on the ‘*Fischer*’ datasets are then validated using independent cohorts (Table 1) [20, 28].

**Figure 1:**
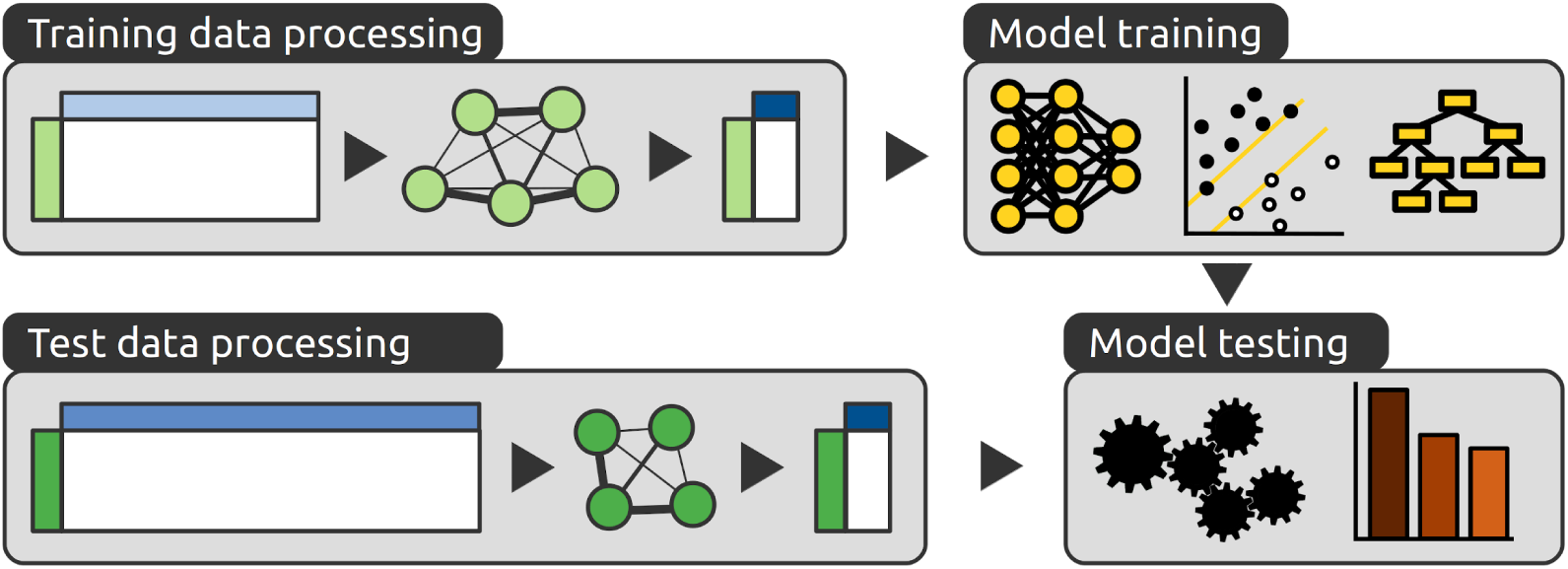
General workflow of the proposed method. Our strategy relies on a topological analysis to perform dimension reduction of both the training (light green) and test data (dark green). Data matrices are transformed into graphs, from which topological features are extracted. Even if the original features (light blues) are different, the topological features extracted from the graphs (dark blue) have the same meaning and are comparable. These features are then used to train and test several models that rely on different learning algorithms (DNN, SVM and RF). These models are compared based on the accuracy of their predictions on the test data.

### Assessment of the topological features

We first compare models that accept different topological features extracted from the ‘*Fischer*’ datasets as input, regardless of the underlying neural network architecture. We have defined nine possible feature sets that can be used as input to the classifiers (Table 2). First, and for each dataset, three feature sets are defined: graph centralities, graph modularities and both combined. Second, we also concatenate the feature sets across the two ‘*Fischer*’ datasets to create three additional feature sets. These feature sets contain between 12 and 101 topological features.

The results of this comparison for the two clinical outcomes can be found in Figure 2. For each feature set, the balanced accuracies over all models (different architectures and replicates) are displayed as a single boxplot. The full list of models and their balanced accuracies is provided in Additional File 3. A first observation is that centrality features are associated with better average performances than modularity features (‘*Death from disease*’, *p* ≤ 1*e*-7; ‘*Disease progression*’, *p* ≤ 1*e*-7). We note that the difference between these average accuracies is modest for ‘*Death from disease*’ (∆_*bACC*_ ∈ [2.4, 3.9]) but more important for ‘*Disease progression*’ (∆_*bACC*_ ∈ [6.7, 8.2]). Combining both types of topological features generally does not improve the average performance.

**Figure 2:**
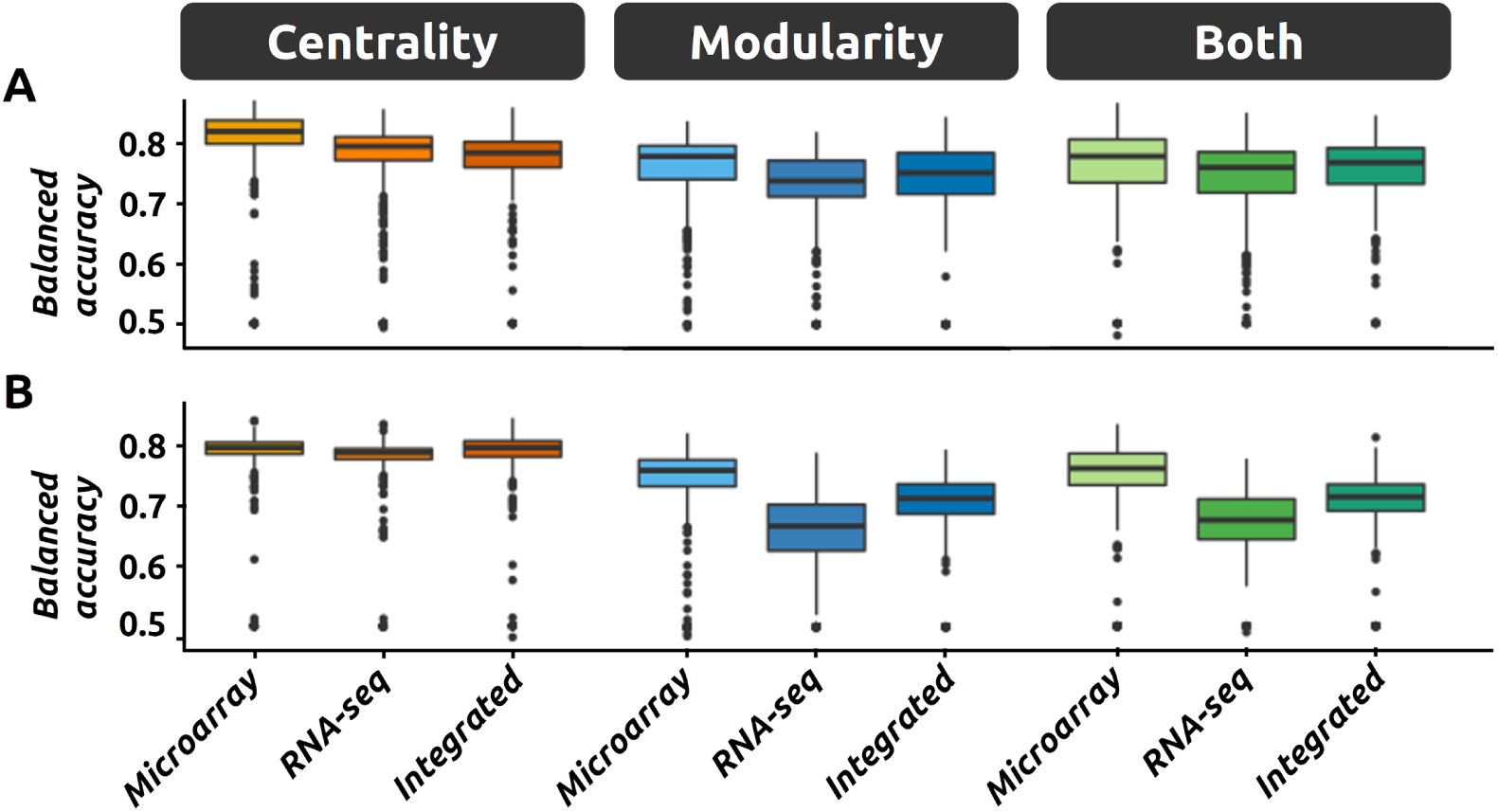
Model performance for different inputs. DNN models relying on different feature sets are compared by reporting their performance on the validation data for ‘*Death from disease*’ (A) and ‘*Disease progression*’ (B). Feature sets are defined by the original data that were used (microarray data, RNA-seq data or the integration of both) and by the topological features considered (centrality, modularity or both). Each single point represents a model. For each feature set, several models are trained by varying the neural network architecture and by performing replicates.

A second observation is that the features extracted from the RNA-seq data are associated with lower average performance than the equivalent features extracted from the microarray data (*p* ≤ 1*e*-7). The differences indicate that once again the effect is not negligible (‘*Death from disease*’, ∆_*bACC*_ ∈ [2.1, 3.6]); ‘*Disease progression*’, ∆_*bACC*_ ∈ [4.4, 6.0]). In addition, the integration of the data across the two expression datasets does not improve the average performance.

### Influence of the DNN architecture

Deep neural networks are feed forward neural networks with several hidden layers, with several nodes each. The network architecture (*i.e.*, layers and nodes) as well as the strategy used to train the network can influence its performance. We have therefore defined 35 possible architectures in total by varying the number of hidden layers and the number of neurons per hidden layer (“Methods”).

We compare the performance of the models relying on these different architectures. The results can be found in Table 3 and Supplementary Figure S1 (Additional File 1). The full list of models and their balanced accuracies is provided in Additional File 3. We can observe a small inverse correlation between the complexity of the architecture and the average performance. Although significant, the average performance of simple models (one hidden layer) is, on average, only marginally better than the average performance of more complex models (at least two hidden layers) (*p* ≤ 1*e*-7, ∆_*bACC*_ ∈ [2, 4]).

**Table 3:** Best performing DNN architectures. One row corresponds to the best model for a given clinical outcome and configuration (from Table 2). The best performance (*i.e.*, balanced accuracy) are displayed in bold for each clinical outcome. ^*a*^ Combined dataset in which the topological features of both ‘*Fischer-M*’ and ‘*Fischer-R*’ are concatenated.

| Configuration | Architecture | Balanced accuracy |
| --- | --- | --- |
| Clinical outcome = ‘*Death from disease*’ |  |  |
| Fischer-M, centralities | [8, 8, 8, 2] | **87.3%** |
| Fischer-M, modularities | [8, 4] | 83.9% |
| Fischer-M, both | [8, 8, 8] | 86.8% |
| Fischer-R, centralities | [8, 8, 8, 4] | 85.8% |
| Fischer-R, modularities | [8, 8, 8, 2] | 82.1% |
| Fischer-R, both | [2, 2, 2, 2] | 85.2% |
| Fischer<sup>a</sup>, centralities | [8, 2, 2] | 86.1% |
| Fischer<sup>a</sup>, modularities | [8, 2, 2] | 84.7% |
| Fischer<sup>a</sup>, both | [8, 8, 4] | 84.7% |
| Clinical outcome = ‘*Disease progression*’ |  |  |
| Fischer-M, centralities | [8, 8, 8, 2] | 84.3% |
| Fischer-M, modularities | [8, 8, 2] | 82.3% |
| Fischer-M, both | [4, 4, 2] | 83.7% |
| Fischer-R, centralities | [8, 8, 4] | 83.7% |
| Fischer-R, modularities | [8, 2, 2] | 79.1% |
| Fischer-R, both | [8, 8, 8, 8] | 77.9% |
| Fischer<sup>a</sup>, centralities | [4, 2, 2, 2] | **84.7%** |
| Fischer<sup>a</sup>, modularities | [8, 8] | 79.6% |
| Fischer<sup>a</sup>, both | [4, 2] | 81.5% |

### Best models

Although the differences in average performance are important, our objective is to identify the best models, regardless of the average performance of any category. In the current section, we therefore report the best models for each feature set and each clinical outcome (summarized in Table 3). In agreement with the global observations, the best model for ‘*Death from disease*’ is based on the centrality features extracted from the microarray data. The best model for ‘*Disease progression*’ relies however on centralities derived from both the microarray and the RNA-seq data (Table 3), even if the corresponding category is not associated with the best average performance. This is consistent with the observation that the variance in performance increases when the number of input features increases, which can produce higher maximum values (Figure 2). We can also observe some level of agreement between the two outcomes of interest. Indeed, the best feature set for ‘*Death from disease*’ is actually the second best for ‘*Disease progression*’. Similarly, the best feature set for ‘*Disease progression*’ is the third best for ‘*Death from disease*’.

Regarding the network architecture, models relying on networks with four hidden layers represent the best models for both ‘*Disease progression*’ and ‘*Death from disease*’ (Table 3). Their respective architectures are still different and the ‘*Disease progression*’ network contains more neurons. However, the second best network for ‘*Disease progression*’ and the best network for ‘*Death from disease*’ share the same architecture (two layers with four neurons each followed by two layers with two neurons each) indicating that this architecture can still perform well in both cases.

### Fine tuning of the hyper-parameters

Based on the previous observations, we have selected the best models for each clinical outcome in order to fine tune their hyper-parameters. The optimization was performed using a simple grid search (“Methods” section). The hyper-parameters we optimized are the learning rate, the optimization strategy and the dropout (included to circumvent over-fitting during training [22]). When considering all models, we can observe that increasing the initial learning rate seems to slightly improve the average performance, although the best models are in fact obtained with a low initial learning rate (Additional File 1, Supplementary Figure S2). The most important impact is observed for the optimization strategies, with the Adam optimizer [15] representing the best strategy, adadelta the less suitable one, with the adagrad variants in between. We observe that the performance is almost invariant to dropout except when it reaches 0.4 where it seems to have a strong negative impact on performance.

When focusing on the best models only, we observe similarities between the two clinical outcomes of interest. Indeed, in both cases, the optimal dropout, optimizer, and learning rate are respectively 0.3, Adam and 1e-3. Notice that for ‘*Death from disease*’, another learning rate value gives exactly the same performance (5e-4). As mentioned above, learning rate has little influence on the average performance. However, for these two specific models, its influence is important and using a non-optimal value results in a drop up to 19% for ‘*Death from disease*’ and 29% for ‘*Disease progression*’. More important, we observe no significant increase in performance after parameter optimization (Table 4), which correlates with the fact that two of the three optimal values actually correspond to the default values that were used before.

**Table 4:** Parameter optimization for all classifiers. One row corresponds to the best model for a given clinical outcome and algorithm. The optimal parameter values are provided (o: optimizer, lr: learning rate, d: dropout, h: sizes of the second and third GEDFN hidden layers, b: batch size, t: SVM kernel type, c: cost, g: gamma, n: number of trees). The gain in balanced accuracy with respect to the models run with default parameters is indicated between parentheses (from Table 3 for DNN). ^*a*^ for GEDFN, the corresponding omics data is used as input instead of the topological features.

| Algorithm | Parameters | Balanced accuracy |
| --- | --- | --- |
| Clinical outcome = ‘*Death from disease*’,<br>Data=*Fischer-M*, centralities |  |  |
| DNN [8,8,8,2] | o=Adam, lr=1e-3, d=0.3 | **87.3%** (+0.0) |
| GEDFN<sup>a</sup> | lr=1e-2, h=[64,16], b=8 | 79.5% (+8.6) |
| SVM | t=RBF, c=64, g=0.25 | 75.4% (+5.9) |
| RF | n=100 | 75.1% (+3.1) |
| Clinical outcome = ‘*Disease progression*’,<br>Data=*Fischer*, centralities |  |  |
| DNN [4,2,2,2] | o=Adam, lr=1e-3, d=0.3 | **84.7%** (+0.0) |
| GEDFN<sup>a</sup> | lr=1e-4, h=[16,4], b=32 | 81.2% (+0.4) |
| SVM | t=RBF, c=16, g=0.0625 | 81.8% (+2.0) |
| RF | n=100 | 78.1% (+3.2) |

Whether we consider the different feature sets or the different network architectures, we also observe that the performance varies across replicates, *i.e.*, models built using the same configuration but different randomization seeds (which are used for sample shuffling and initialization of the weights and biases). This seems to indicate that better models might also be produced simply by running more replicates. We tested this hypothesis by running more replicates of the best configurations (*i.e.*, increasing the number of replicates from 10 to 100). However, we report no improvement of these models with 90 additional replicates (Additional File 3).

### Comparison to other modeling strategies

We then compare the DNN classifiers to other classifiers relying on different learning algorithms (SVM and RF). These algorithms have previously demonstrated their effectiveness to solve the same classification task on the ‘*Fischer*’ dataset, albeit using a different patient stratification [37, 25]. For a fair comparison, all classifiers are input the same features and are trained and tested using the same samples. Optimal performance is obtained via a grid search over the parameter space (“Methods” section). The results are summarized in Table 4. We observe that the DNN classifiers outperform both the SVM and RF classifiers for both outcomes (‘*Death from disease*’, ∆_*bACC*_ ∈ [11.9, 12.2]); ‘*Disease progression*’, ∆_*bACC*_ ∈ [2.9, 6.6]).

We also compare our strategy to GEDFN, an approach based on a neural network which requires a feature graph to enforce sparsity of the connections between the input and the first hidden layers. Unlike the other models, GEDFN models only accept omics data as input (*i.e.*, original features). They are also optimized using a simple grid search. The results are summarized in Table 4. We can observe that the GEDFN models perform better than the SVM and RF models for ‘*Death from disease*’. For ‘*Disease progression*’, the GEDFN and SVM models are on par, and both superior to RF models. For both clinical outcomes, the GEDFN models remain however less accurate than the DNN models that use topological features. (‘*Death from disease*’, ∆_*bACC*_ = 7.8); ‘*Disease progression*’, ∆_*bACC*_ = 3.5)

### Validation with independent datasets

In a last set of experiments, we tested our models using independent datasets. First, we use the ‘*Fischer-M*’ dataset to validate models built using the ‘*Fischer-R*’ dataset and vice-versa. Then, we also make use of two fully independent datasets, ‘*Maris*’ and ‘*Versteeg*’ as validation datasets for all models trained with any of the ‘*Fischer*’ datasets. We compare the performance on these independent datasets to the reference performance (obtained when the same dataset is used for both training and testing).

The results are summarized in Table 5. When one of the ‘*Fischer*’ dataset is used for training and the other dataset for testing, we can, in general, observe a small decrease in performance with respect to the reference (DNN, ∆_*bACC*_ ∈ [3.7., 7.3]; SVM, ∆_*bACC*_ ∈ [*−*9.4, 21.9]; RF, ∆_*bACC*_ ∈ [*−*1.7, 8.3]). For SVM and RF models, there is sometimes an increased performance (negative ∆_*bACC*_).

**Table 5:** External validation results. Models are trained using one of the ‘*Fischer*’ datasets and then tested using either the other ‘*Fischer*’ dataset or another independent dataset (‘*Maris*’ and ‘*Versteeg*’). The ‘*Maris*’ and ‘*Versteeg*’ datasets are too small to be used for both training and therefore are only used for validation. Rows in italics represent reference models (training and testing extracted from the same datasets).

| Datasets |  | Balanced accuracy |  |  |
| --- | --- | --- | --- | --- |
| Training | Test | DNN | SVM | RF |
| Clinical outcome = ‘*Death from disease*’,<br>Data = centralities |  |  |  |  |
| *Fischer-M* | *Fischer-M* | *87.3%* | *75.4%* | *75.1%* |
|  | *Fischer-R* | **82.1%** | 53.5% | 66.8% |
|  | *Maris* | 53.1% | **54.3%** | 50.0% |
|  | *Versteeg* | **75.0%** | 53.3% | 67.5% |
| *Fischer-R* | *Fischer-R* | *85.8%* | *66.0%* | *62.4%* |
|  | *Fischer-M* | **81.5%** | 75.4% | 61.2% |
|  | *Maris* | **56.2%** | 49.7% | 50.0% |
|  | *Versteeg* | **70.8%** | 68.3% | 67.5% |
| Clinical outcome = ‘*Disease progression*’,<br>Data = centralities |  |  |  |  |
| *Fischer-M* | *Fischer-M* | *84.3%* | *83.7%* | *80.0%* |
|  | *Fischer-R* | **77.0%** | 75.2% | 71.8% |
|  | *Maris* | **67.5%** | 66.0% | 53.8% |
|  | *Versteeg* | 78.1% | **82.4%** | 78.1% |
| *Fischer-R* | *Fischer-R* | *83.7%* | *81.0%* | *73.3%* |
|  | *Fischer-M* | **80.0%** | 76.8% | 75.0% |
|  | *Maris* | **67.5%** | 58.8% | 58.8% |
|  | *Versteeg* | **80.1%** | 77.2% | 73.9% |

When considering the fully independent datasets, we observe two different behaviors. For the ‘*Maris*’ dataset, the performance ranges from random-like (DNN, 53% and 56%) to average (DNN, 68%) for ‘*Death from disease*’ and ‘*Disease progression*’ respectively. Similar results are obtained for SVM and RF models (between 50% and 66%). Altogether, these results indicate that none of the models is able to classify the samples of this dataset. However, for the ‘*Versteeg*’ dataset, and for both clinical outcomes, the models are more accurate (DNN, from 71% to 80%), in the range of the state of the art for neuroblastoma. A similar trend is observed for the SVM and RF models, although the DNN models seem superior in most cases. The drop in performance for *Versteeg*’ (with respect to the reference models) is within the same range than for ‘*Fischer*’ (DNN, ∆_*bACC*_ ∈ [3.6., 15.0]; SVM, ∆_*bACC*_ ∈ [−2.3, 22.1]; RF, ∆_*bACC*_ ∈ [−5.1, 7.6]). For both ‘*Maris*’ and ‘*Versteeg*’ datasets, it is difficult to appreciate the classification accuracies in the absence of reference models, due to the small number of samples available for these two cohorts (less than 100).

## Discussion

We evaluate several strategies to build models that use expression profiles of patients as input to classify patients according to their clinical outcomes. We propose to tackle the “small n large p” problem, frequently associated with such omics datasets, via a graph-based dimension reduction method. We have applied our approach to four neuroblastoma datasets to create and optimize models based on their classification accuracy.

We first investigate the usefulness of different sets of topological features by measuring the performance of classification models using different inputs. We observe that centrality features are associated with better average performances than modularity features. We also note that the features extracted from the RNA-seq data are associated with lower performance than the equivalent features extracted from the microarray data. Both seems to contradict our previous study of the same classification problem, in which we reported no statistical difference between models built from both sets [25]. It is important to notice however that the learning algorithms and the data stratification are different between the two studies, which might explain this discrepancy. In addition, the accuracies reported here are often greater than the values reported previously, but not always by the same margin, which creates differences that were not apparent before. We also observe that the difference is mostly driven by the weak performance of models relying on the modularity features extracted from the ‘*Fischer-R*’ dataset. This suggests that although the individual RNA-sequencing features do correlate with clinical outcomes, their integration produces modules whose correlation is lower (in comparison to microarray data). This corroborates a recent observation that deriving meaningful modules from WGCNA co-expression graphs can be rather challenging [**?**].

We observe that the combined feature sets are not associated with any improvement upon the individual feature sets. This indicates that both sets might actually measure the same topological signal, which is in line with our previous observations [25]. Similarly, the integration of the data across the two expression datasets does not improve the average performance. This was rather expected since the two datasets measure the same biological signal (*i.e.*, gene expression) albeit through the use of different technologies.

Neural networks are known to be rather challenging to optimize, and a small variation in one parameter can have dramatic consequences, especially when the set of instances is rather limited. We indeed observe important variations in performance within the categories we have defined (*e.g.*, models using only centralities or four layer DNN models) as illustrated in Figure 2 and Supplementary Figures S1 and S2.

The parameters with the greatest influence on performance are the optimization strategy (Adam really seems superior in our case) and the dropout (it should be below 0.4). In the latter case, it is not surprising that ignoring at least 40% of the nodes can have a huge impact on networks that have less than 100 input nodes and at best 8 nodes per hidden layer.

Regarding the network architecture, models relying on four layer networks perform the best for both clinical outcomes (Table 3). This is in agreement with previous studies that have reported that such relatively small networks (*i.e.*, with three or four layers) can efficiently predict clinical outcomes of kidney cancer patients [17] or can capture relevant features for survival analyses of a neuroblastoma cohort [12].

Even if there are differences, as highlighted above, the optimal models and parameters are surprisingly similar for both clinical outcomes. This is true for the input data, the network architecture and the optimal values of the hyper-parameters. We also note, however, that this might be due to the underlying correlation between the two clinical outcomes since almost all patients who died from the disease have experienced progression of the disease.

When applied on the ‘*Fischer*’ datasets, the DNN classifiers outperform both SVM and RF classifiers for both outcomes. The gain in performance is modest for ‘*Disease progression*’ but rather large for ‘*Death from disease*’, which was previously considered as the hardest outcome to predict among the two [37].

We also compare our neural networks fed with graph topological features (DNN) to neural networks fed with expression profiles directly (GEDFN). We notice that the GEDFN models perform at least as good as the SVM and RF models, but also that they remain less accurate than the DNN models. Altogether these observations support the idea that deep neural networks could indeed be more effective than traditional SVM and RF models. In addition, it seems that coupling such deep neural networks with a graph-based topological analysis can give even more accurate models.

Last, we validate the models using independent datasets. The hypothesis of these experiments is that the topological features we derived from the omics data are independent of the technology used in the first place and can therefore enable better generalization. As long as a graph of patients (PSN) can be created, it will be possible to derive topological features even if microarrays have been used in one study and sequencing in another study (or any other biomedical data for that matter). We therefore hypothesize that a model trained using one cohort might be tested using another cohort, especially when this second cohort is too small to be used to train another model by itself.

We start by comparing the two ‘*Fischer*’ datasets. As expected, we observe a small decrease in performance in most cases when applying the models on the ‘*Fischer*’ dataset that was not used for training. Surprisingly, for SVM and RF, the performance for the independent datasets is sometimes better than the reference performance. However, this happens only when the reference performance is moderate at best (*i.e.*, *bACC <* 75%). For DNN models, the performance on the independent datasets is still reasonable (at least 81.5% and 77% for ‘*Death from disease*’ and ‘*Disease progression*’ respectively) and sometimes even better than reference SVM and RF models (in six of the eight comparisons, Table 5).

We then include two additional datasets that are too small to be used to train classification models (‘*Maris*’ and ‘*Versteeg*’ datasets). Similarly to above, we note that, in most cases, the DNN models are more accurate than the corresponding SVM and RF models, especially for the ‘*Death from disease*’ outcome. Regarding the poor overall performance on the ‘*Maris*’ dataset, we observe that it is the oldest of the datasets, associated with one of the first human high-throughput microarray platform (HG-U95A), that contains less probes than there are human genes (Table 2). In addition, we note that the median patient follow-up for this dataset was 2.3 years, which, according to the authors of the original publication, was too short to allow them to study the relationship between expression profiles and clinical outcome, in particular patient survival [28] (page 6052). In contrast, the median patient follow-up for the ‘*Versteeg*’ dataset was 12.5 years, which allows for a more accurate measure of long term clinical outcomes. Altogether, these reasons might explain why the performance remains poor for the ‘*Maris*’ dataset (especially for ‘*Death from disease*’) in contrast to the other datasets.

## Conclusion

We propose a graph-based method to extract features from patient derived omics data. These topological features are then used as input to a deep neural network that can classify patients according to their clinical outcome. Our models can handle typical omics datasets (with small *n* and large *p*) first, by reducing the number of features (through extraction of topological features) and second, by fine tuning the deep neural networks and their parameters.

By applying our strategy to four neuroblastoma datasets, we observe that our models make more accurate predictions than models based on other algorithms or different strategies. This indicates that the deep neural networks are indeed capturing complex features in the data that other machine learning strategies might not. In addition, we also demonstrate that our graph-based feature extraction method allows to validate the trained models using external datasets, even when the original features are different.

Additional studies are however needed to explore the properties of these topological features and their usefulness when coupled to deep learning predictors. In particular, applications to other data types (beside gene expression data) and other genetic disorders (beside neuroblastoma) are necessary to validate the global utility of the proposed approach. Moreover, other modeling strategies that integrate graphs (and their topology) into the learning process, such as graph-based CNN [8, 16] would need to be explored as well.

## Abbreviations

bACC: balanced accuracy
CNN: Convolutional Neural Network
DNN: Deep Neural Network
GEO: Gene Expression Omnibus
PPIN: Protein-Protein Interaction Network
PSN: Patient Similarity Networks
ReLU: Rectified Linear Unit
RBF: Radial Basis Function
RF: Random Forest
RNA: RiboNucleic Acid
SBM: Stochastic Block Model
SVM: Support Vector Machine
TCGA: The Cancer Genome Atlas
WGCNA: Weighted Correlation Network Analysis

## Availability of data and materials

The code associated to this paper can be found at https://gitlab.com/biomodlih/SingalunDeep The input data for the classifiers (*i.e.*, topological features) are available on Zenodo (10.5281/zenodo.3357673, version 1.0.0). These data are derived from the publicly available datasets we use (available on GEO and R2 – see methods).

## Competing interests

The authors declare that they have no competing interests.

## Funding

Project supported by the Fonds National de la Recherche (FNR), Luxembourg (SINGALUN project). This research was also partially supported by Tier-2 grant MOE2016-T2-1-029 by the Ministry of Education, Singapore. The funders had no role in study design, data collection and analysis, decision to publish, or preparation of the manuscript.

## Author’s contributions

All authors have developed the strategy. LT has implemented the method and applied it to the neuroblastoma datasets. All authors have analyzed the results. LT wrote an initial draft of the manuscript. All authors have revised the manuscript. All authors read and approved the final manuscript.

## Acknowledgements

We thank the Fischer, Maris and Veersteeg laboratories for sharing their neuroblastoma data. In particular, we thank Dr John M. Maris and Dr Alvin Farrel for helping us with the clinical data of their cohort. We thank Dr Liyanaarachchi Lekamalage Chamara Kasun for helpful discussion about the DNN models. We thank Tony Kaoma and Dr Petr V. Nazarov for helpful discussions regarding the model comparison. We thank Dr Enrico Glaab and Dr Rama Kaalia for their support during the project.

## Notes

#### Summary of Updates

The "Results and discussion" section has been split into two sections "Results" and "Discussion". The text of these two sections has been accommodated accordingly.

https://gitlab.com/biomodlih/SingalunDeep

https://zenodo.org/record/3357674

## References

[1] Gene expression omnibus. https://www.ncbi.nlm.nih.gov/geo/.

[2] R2: Genomics analysis and visualization platform. https://hgserver1.amc.nl/cgi-bin/r2/main.cgi.

[3] A. C. Berger, A. Korkut, R. S. Kanchi, A. M. Hegde, W. Lenoir, W. Liu, Y. Liu, H. Fan, H. Shen, V. Ravikumar, A. Rao, A. Schultz, X. Li, P. Sumazin, C. Williams, P. Mestdagh, P. H. Gunaratne, C. Yau, and R. Bowlby. A comprehensive pan-cancer molecular study of gynecologic and breast cancers. Cancer Cell, 33(4):690–705.e9.

[4] P. Calvas, L. Jamot, J. Weinbach, N. Chassaing, and T. RaDiCo Team. The RaDiCo AC-OEIL: a french rare disease cohort dedicated to ocular developmental anomalies in children. Acta Ophthalmologica, 95.

[5] J. Das and H. Yu. HINT: High-quality protein interactomes and their applications in understanding human disease. BMC Systems Biology, 6:92.

[6] J. N. De Roach, T. L. McLaren, R. L. Paterson, E. C. O’Brien, L. Hoffmann, D. A. Mackey, A. W. Hewitt, and T. M. Lamey. Establishment and evolution of the australian inherited retinal disease register and DNA bank. Clin. Experiment. Ophthalmol., 41(5):476–483.

[7] A. Decelle, F. Krzakala, C. Moore, and L. Zdeborová. Asymptotic analysis of the stochastic block model for modular networks and its algorithmic applications. Phys. Rev. E, 84(6):066106.

[8] M. Defferrard, X. Bresson, and P. Vandergheynst. Convolutional neural networks on graphs with fast localized spectral filtering. arXiv:1606.09375 [cs, stat].

[9] J. Dutkowski and T. Ideker. Protein networks as logic functions in development and cancer. PLoS Comput. Biol., 7(9):e1002180.

[10] H. Feng, P. Jin, and H. Wu. Disease prediction by cell-free DNA methylation. Brief. Bioinformatics.

[11] H. V. Firth, S. M. Richards, A. P. Bevan, S. Clayton, M. Corpas, D. Rajan, S. V. Vooren, Y. Moreau, R. M. Pettett, and N. P. Carter. DECIPHER: Database of chromosomal imbalance and phenotype in humans using ensembl resources. Am J Hum Genet, 84(4):524–533.

[12] M. Francescatto, M. Chierici, S. Rezvan Dezfooli, A. Zandonà, G. Jurman, and C. Furlanello. Multi-omics integration for neuroblastoma clinical endpoint prediction. Biol. Direct, 13(1):5.

[13] R. G. Jayasinghe, S. Cao, Q. Gao, M. C. Wendl, N. S. Vo, S. M. Reynolds, Y. Zhao, H. Climente-González, S. Chai, F. Wang, R. Varghese, M. Huang, W.-W. Liang, M. A. Wyczalkowski, S. Sengupta, Z. Li, S. H. Payne, D. Fenyö, J. H. Miner, and M. J. Walter. Systematic analysis of splice-site-creating mutations in cancer. Cell Reports, 23(1):270–281.e3.

[14] J. Katzman, U. Shaham, J. Bates, A. Cloninger, T. Jiang, and Y. Kluger. DeepSurv: Personalized treatment recommender system using a cox proportional hazards deep neural network. BMC Medical Research Methodology, 18(1).

[15] D. P. Kingma and J. Ba. Adam: A method for stochastic optimization. arXiv:1412.6980 [cs].

[16] T. N. Kipf and M. Welling. Semi-supervised classification with graph convolutional networks. arXiv:1609.02907 [cs, stat].

[17] Y. Kong and T. Yu. A graph-embedded deep feedforward network for disease outcome classification and feature selection using gene expression data. Bioinformatics.

[18] M. B. Kursa. Robustness of random forest-based gene selection methods. BMC Bioinformatics, 15:8.

[19] T. Liloglou, N. G. Bediaga, B. R. B. Brown, J. K. Field, and M. P. A. Davies. Epigenetic biomarkers in lung cancer. Cancer Lett., 342(2):200–212.

[20] J. J. Molenaar, J. Koster, D. A. Zwijnenburg, P. van Sluis, L. J. Valentijn, I. van der Ploeg, M. Hamdi, J. van Nes, B. A. Westerman, J. van Arkel, M. E. Ebus, F. Haneveld, A. Lakeman, L. Schild, P. Molenaar, P. Stroeken, M. M. van Noesel, I. Ora, E. E. Santo, H. N. Caron, E. M. Westerhout, and R. Versteeg. Sequencing of neuroblastoma identifies chromothripsis and defects in neuritogenesis genes. Nature, 483(7391):589–593.

[21] D. O. Mook-Kanamori, M. M. E.-D. Selim, A. H. Takiddin, H. Al-Homsi, K. A. S. Al-Mahmoud, A. Al-Obaidli, M. A. Zirie, J. Rowe, N. A. Yousri, E. D. Karoly, T. Kocher, W. Sekkal Gherbi, O. M. Chidiac, M. J. Mook-Kanamori, S. Abdul Kader, W. A. Al Muftah, C. McKeon, and K. Suhre. 1,5-anhydroglucitol in saliva is a noninvasive marker of short-term glycemic control. J. Clin. Endocrinol. Metab., 99(3):E479–483.

[22] N. Srivastava, G. Hinton, A. Krizhevsky, I. Sutskever, and R. Salakhutdinov. Dropout: A simple way to prevent neural networks from overfitting. Journal of Machine Learning Research, 15:1929–1958.

[23] K. Suhre, M. Arnold, A. M. Bhagwat, R. J. Cotton, R. Engelke, J. Raffler, H. Sarwath, G. Thareja, A. Wahl, R. K. DeLisle, L. Gold, M. Pezer, G. Lauc, M. A. El-Din Selim, D. O. Mook-Kanamori, E. K. Al-Dous, Y. A. Mohamoud, J. Malek, K. Strauch, H. Grallert, A. Peters, G. Kastenmüller, C. Gieger, and J. Graumann. Connecting genetic risk to disease end points through the human blood plasma proteome. Nat Commun, 8:14357.

[24] The Cancer Genome Atlas Research Network. Comprehensive, Integrative Genomic Analysis of Diffuse Lower-Grade Gliomas. New England Journal of Medicine, 372(26):2481–2498, June 2015.

[25] L.-C. Tranchevent, P. V. Nazarov, T. Kaoma, G. P. Schmartz, A. Muller, S.-Y. Kim, J. C. Rajapakse, and F. Azuaje. Predicting clinical outcome of neuroblastoma patients using an integrative network-based approach. Biol. Direct, 13(1):12.

[26] B. Wang, A. M. Mezlini, F. Demir, M. Fiume, Z. Tu, M. Brudno, B. Haibe-Kains, and A. Goldenberg. Similarity network fusion for aggregating data types on a genomic scale. Nature Methods, 11(3):333–337, Jan. 2014.

[27] C. Wang, B. Gong, P. R. Bushel, J. Thierry-Mieg, D. Thierry-Mieg, J. Xu, H. Fang, H. Hong, J. Shen, Z. Su, J. Meehan, X. Li, L. Yang, H. Li, P. P. Łabaj, D. P. Kreil, D. Megherbi, S. Gaj, F. Caiment, J. van Delft, J. Kleinjans, A. Scherer, V. Devanarayan, J. Wang, Y. Yang, H.-R. Qian, L. J. Lancashire, M. Bessarabova, Y. Nikolsky, C. Furlanello, M. Chierici, D. Albanese, G. Jurman, S. Riccadonna, M. Filosi, R. Visintainer, K. K. Zhang, J. Li, J.-H. Hsieh, D. L. Svoboda, J. C. Fuscoe, Y. Deng, L. Shi, R. S. Paules, S. S. Auerbach, and W. Tong. The concordance between RNA-seq and microarray data depends on chemical treatment and transcript abundance. Nat. Biotechnol., 32(9):926–932.

[28] Q. Wang, S. Diskin, E. Rappaport, E. Attiyeh, Y. Mosse, D. Shue, E. Seiser, J. Jagannathan, S. Shusterman, M. Bansal, D. Khazi, C. Winter, E. Okawa, G. Grant, A. Cnaan, H. Zhao, N.-K. Cheung, W. Gerald, W. London, K. K. Matthay, G. M. Brodeur, and J. M. Maris. Integrative genomics identifies distinct molecular classes of neuroblastoma and shows that multiple genes are targeted by regional alterations in DNA copy number. Cancer Res., 66(12):6050–6062.

[29] Z. Wang, B. Yang, M. Zhang, W. Guo, Z. Wu, Y. Wang, L. Jia, S. Li, S. J. Caesar-Johnson, J. A. Demchok, I. Felau, M. Kasapi, M. L. Ferguson, C. M. Hutter, H. J. Sofia, R. Tarnuzzer, Z. Wang, L. Yang, J. C. Zenklusen, and J. Zhang. lncRNA epigenetic landscape analysis identifies EPIC1 as an oncogenic lncRNA that interacts with MYC and promotes cell-cycle progression in cancer. Cancer Cell, 33(4):706–720.e9.

[30] G. P. Way, F. Sanchez-Vega, K. La, J. Armenia, W. K. Chatila, A. Luna, C. Sander, A. D. Cherniack, M. Mina, G. Ciriello, N. Schultz, Cancer Genome Atlas Research Network, Y. Sanchez, and C. S. Greene. Machine learning detects pan-cancer ras pathway activation in the cancer genome atlas. Cell Rep, 23(1):172–180.e3.

[31] B. Xiao, W. Zhang, L. Chen, J. Hang, L. Wang, R. Zhang, Y. Liao, J. Chen, Q. Ma, Z. Sun, and L. Li. Analysis of the miRNA-mRNA-lncRNA network in human estrogen receptor-positive and estrogen receptor-negative breast cancer based on TCGA data. Gene, 658:28–35.

[32] S. Yousefi, F. Amrollahi, M. Amgad, C. Dong, J. E. Lewis, C. Song, D. A. Gutman, S. H. Halani, J. E. Velazquez Vega, D. J. Brat, and L. A. D. Cooper. Predicting clinical outcomes from large scale cancer genomic profiles with deep survival models. Sci Rep, 7(1):11707.

[33] S. Yousefi, C. Song, N. Nauata, and L. Cooper. Learning genomic representations to predict clinical outcomes in cancer. arXiv:1609.08663 [cs].

[34] K.-H. Yu, D. A. Levine, H. Zhang, D. W. Chan, Z. Zhang, and M. Snyder. Predicting ovarian cancer patients’ clinical response to platinum-based chemotherapy by their tumor proteomic signatures. J. Proteome Res., 15(8):2455–2465.

[35] D. R. Zerbino, P. Achuthan, W. Akanni, M. R. Amode, D. Barrell, J. Bhai, K. Billis, C. Cummins, A. Gall, C. G. Girón, L. Gil, L. Gordon, L. Haggerty, E. Haskell, T. Hourlier, O. G. Izuogu, and S. H. Janacek. Ensembl 2018. Nucleic Acids Res, 46:D754–D761.

[36] B. Zhang and S. Horvath. A general framework for weighted gene co-expression network analysis. Statistical Applications in Genetics and Molecular Biology, 4:Article17, 2005.

[37] W. Zhang, Y. Yu, F. Hertwig, J. Thierry-Mieg, W. Zhang, D. Thierry-Mieg, J. Wang, C. Furlanello, V. Devanarayan, J. Cheng, Y. Deng, B. Hero, H. Hong, M. Jia, L. Li, S. M. Lin, Y. Nikolsky, A. Oberthuer, T. Qing, and Z. Su. Comparison of RNA-seq and microarray-based models for clinical endpoint prediction. Genome Biology, 16(1), Dec. 2015.

