## Supplementary material for "A deep neural network approach to predicting clinical outcomes of neuroblastoma patients"

### Additional file 1 for "A deep neural network approach to predict clinical outcomes of Neuroblastoma patients"

#### List of Figures

S1 Performance of the DNN models with various architectures. . . . . . . . . . . . . . . 1
S2 Detailed performance of the models trained with different parameters. . . . . . . . 2

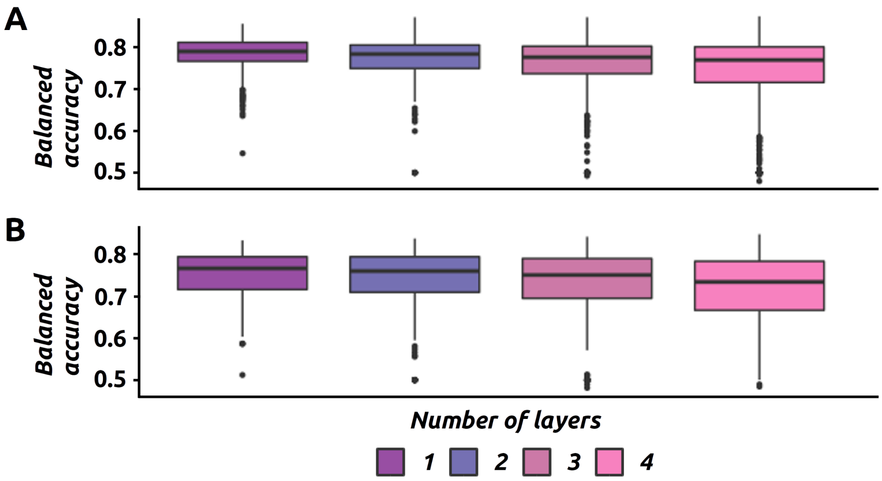


Supplementary Figure S1: The performance (*i.e.*, balanced accuracy) of DNN models with different architectures (*i.e.*, number of layers) for '*Death from disease*' (A) and '*Disease progression*' (B).

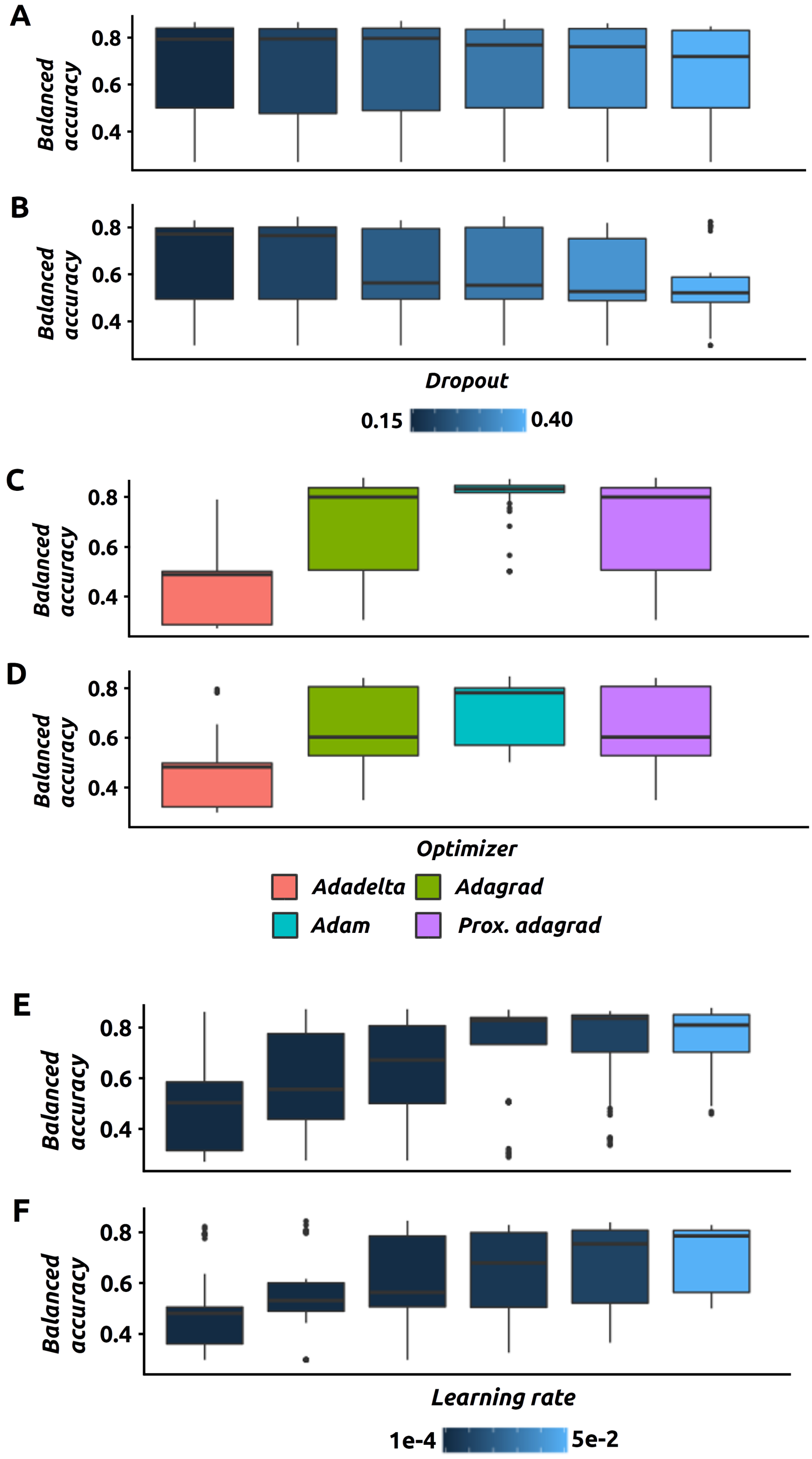


Supplementary Figure S2: The performance (*i.e.*, balanced accuracy) of DNN models in function of hyper-parameter such as dropout (A and B), optimizing strategy (C and D) and learning rate (E and F). Results are displayed for '*Death from disease*' (A, C and E) and '*Disease progression*' (B, D and F).

| Train_part1 | | | | | Train_part2 | | | | | Eval | | | | | Valid | | | | |
|---|---|---|---|---|---|---|---|---|---|---|---|---|---|---|---|---|---|---|---|
| **Patient_id** | **Gender** | **High-risk** | **rogressio** | **death from disease** | **Patient_id** | **Gender** | **High-risk** | **rogressio** | **death from disease** | **Patient_id** | **Gender** | **High-risk** | **rogressio** | **death from disease** | **Patient_id** | **Gender** | **High-risk** | **rogressio** | **death from disease** |
| NB002 | 0 | 1 | 1 | 1 | NB004 | 1 | 1 | 0 | 0 | NB001 | 0 | 1 | 1 | 1 | NB003 | 1 | 1 | 1 | 1 |
| NB006 | 0 | 0 | 0 | 0 | NB008 | 0 | 1 | 0 | 0 | NB005 | 1 | 1 | 1 | 1 | NB007 | 0 | 1 | 1 | 0 |
| NB010 | 0 | 1 | 1 | 1 | NB012 | 0 | 0 | 0 | 0 | NB009 | 1 | 1 | 0 | 0 | NB011 | 0 | 0 | 0 | 0 |
| NB014 | 1 | 0 | 0 | 0 | NB016 | 1 | 0 | 0 | 0 | NB013 | 0 | 1 | 1 | 1 | NB015 | 0 | 1 | 1 | 0 |
| NB018 | 0 | 0 | 0 | 0 | NB020 | 0 | 0 | 0 | 0 | NB017 | 0 | 0 | 0 | 0 | NB019 | 1 | 0 | 0 | 0 |
| NB022 | 0 | 0 | 0 | 0 | NB024 | 1 | 0 | 0 | 0 | NB021 | 1 | 0 | 0 | 0 | NB023 | 0 | 0 | 0 | 0 |
| NB026 | 1 | 0 | 0 | 0 | NB028 | 1 | 0 | 0 | 0 | NB025 | 0 | 0 | 0 | 0 | NB027 | 1 | 0 | 0 | 0 |
| NB030 | 0 | 0 | 0 | 0 | NB032 | 1 | 0 | 0 | 0 | NB029 | 0 | 0 | 0 | 0 | NB031 | 1 | 0 | 0 | 0 |
| NB034 | 1 | 0 | 0 | 0 | NB036 | 0 | 0 | 0 | 0 | NB033 | 1 | 0 | 0 | 0 | NB035 | 0 | 0 | 0 | 0 |
| NB038 | 1 | 0 | 0 | 0 | NB040 | 0 | 0 | 0 | 0 | NB037 | 1 | 0 | 0 | 0 | NB039 | 1 | 0 | 0 | 0 |
| NB042 | 1 | 0 | 0 | 0 | NB044 | 1 | 0 | 0 | 0 | NB041 | 1 | 0 | 0 | 0 | NB043 | 0 | 0 | 0 | 0 |
| NB046 | 1 | 0 | 0 | 0 | NB048 | 0 | 0 | 0 | 0 | NB045 | 1 | 0 | 0 | 0 | NB047 | 1 | 0 | 0 | 0 |
| NB050 | 1 | 0 | 0 | 0 | NB052 | 1 | 0 | 0 | 0 | NB049 | 1 | 0 | 0 | 0 | NB051 | 0 | 0 | 0 | 0 |
| NB054 | 1 | 0 | 0 | 0 | NB056 | 0 | 0 | 0 | 0 | NB053 | 0 | 0 | 0 | 0 | NB055 | 1 | 0 | 0 | 0 |
| NB058 | 1 | 0 | 0 | 0 | NB060 | 0 | 0 | 0 | 0 | NB057 | 0 | 0 | 0 | 0 | NB059 | 1 | 0 | 0 | 0 |
| NB062 | 1 | 0 | 0 | 0 | NB064 | 0 | 0 | 0 | 0 | NB061 | 0 | 0 | 0 | 0 | NB063 | 1 | 0 | 1 | 0 |
| NB066 | 1 | 0 | 1 | 0 | NB068 | 1 | 0 | 0 | 0 | NB065 | 0 | 0 | 0 | 0 | NB067 | 0 | 0 | 0 | 0 |
| NB070 | 1 | 0 | 0 | 0 | NB072 | 0 | 0 | 0 | 0 | NB069 | 1 | 0 | 1 | 1 | NB071 | 1 | 0 | 0 | 0 |
| NB074 | 0 | 0 | 1 | 0 | NB076 | 0 | 0 | 0 | 0 | NB073 | 1 | 0 | 0 | 0 | NB075 | 1 | 0 | 0 | 0 |
| NB078 | 0 | 0 | 0 | 0 | NB080 | 1 | 1 | 1 | 1 | NB077 | 0 | 0 | 0 | 0 | NB079 | 0 | 0 | 0 | 0 |
| NB082 | 0 | 1 | 0 | 0 | NB084 | 1 | 0 | 0 | 0 | NB081 | 0 | 1 | 0 | 0 | NB083 | 0 | 1 | 0 | 0 |
| NB086 | 1 | 0 | 1 | 0 | NB088 | 1 | 0 | 0 | 0 | NB085 | 1 | 0 | 0 | 0 | NB087 | 0 | 1 | 1 | 0 |
| NB090 | 1 | 1 | 1 | 1 | NB092 | 0 | 1 | 1 | 1 | NB089 | 0 | 0 | 0 | 0 | NB091 | 0 | 1 | 1 | 0 |
| NB094 | 0 | 1 | 1 | 1 | NB096 | 1 | 0 | 0 | 0 | NB093 | 0 | 0 | 0 | 0 | NB095 | 0 | 0 | 0 | 0 |
| NB098 | 0 | 0 | 0 | 0 | NB100 | 0 | 0 | 0 | 0 | NB097 | 1 | 0 | 0 | 0 | NB099 | 0 | 0 | 0 | 0 |
| NB102 | 0 | 0 | 0 | 0 | NB104 | 0 | 0 | 0 | 0 | NB101 | 0 | 0 | 0 | 0 | NB103 | 1 | 0 | 1 | 0 |
| NB106 | 0 | 0 | 1 | 0 | NB108 | 0 | 0 | 0 | 0 | NB105 | 1 | 0 | 0 | 0 | NB107 | 1 | 0 | 0 | 0 |
| NB110 | 0 | 0 | 0 | 0 | NB112 | 1 | 0 | 0 | 0 | NB109 | 0 | 0 | 1 | 0 | NB111 | 1 | 0 | 0 | 0 |
| NB114 | 0 | 0 | 0 | 0 | NB116 | 1 | 0 | 0 | 0 | NB113 | 0 | 1 | 1 | 0 | NB115 | 0 | 0 | 0 | 0 |
| NB118 | 0 | 0 | 0 | 0 | NB120 | 0 | 0 | 0 | 0 | NB117 | 1 | 0 | 1 | 0 | NB119 | 1 | 0 | 1 | 1 |
| NB126 | 1 | 0 | 0 | 0 | NB122 | 1 | 0 | 1 | 1 | NB121 | 0 | 0 | 0 | 0 | NB123 | 0 | 0 | 0 | 0 |
| NB130 | 0 | 0 | 1 | 0 | NB124 | 0 | 0 | 1 | 0 | NB125 | 1 | 0 | 0 | 0 | NB127 | 1 | 0 | 0 | 0 |
| NB134 | 0 | 0 | 0 | 0 | NB128 | 0 | 0 | 0 | 0 | NB129 | 0 | 0 | 0 | 0 | NB131 | 0 | 0 | 0 | 0 |
| NB138 | 0 | 1 | 0 | 0 | NB132 | 0 | 0 | 0 | 0 | NB133 | 1 | 0 | 1 | 0 | NB135 | 0 | 0 | 0 | 0 |
| NB142 | 1 | 0 | 0 | 0 | NB136 | 0 | 1 | 1 | 1 | NB137 | 0 | 1 | 1 | 1 | NB139 | 1 | 0 | 0 | 0 |
| NB146 | 0 | 0 | 0 | 0 | NB140 | 1 | 1 | 1 | 1 | NB141 | 0 | 0 | 0 | 0 | NB143 | 1 | 1 | 0 | 0 |
| NB150 | 1 | 0 | 0 | 0 | NB144 | 0 | 1 | 0 | 0 | NB145 | 0 | 0 | 0 | 0 | NB147 | 0 | 0 | 0 | 0 |
| NB154 | 0 | 0 | 0 | 0 | NB148 | 0 | 0 | 0 | 0 | NB149 | 1 | 0 | 1 | 1 | NB151 | 1 | 0 | 0 | 0 |
| NB158 | 1 | 0 | 0 | 0 | NB152 | 0 | 0 | 0 | 0 | NB153 | 1 | 0 | 0 | 0 | NB155 | 0 | 0 | 0 | 0 |
| NB162 | 1 | 0 | 0 | 0 | NB156 | 0 | 0 | 0 | 0 | NB157 | 0 | 0 | 0 | 0 | NB159 | 1 | 0 | 0 | 0 |
| NB166 | 0 | 0 | 0 | 0 | NB160 | 0 | 0 | 0 | 0 | NB161 | 1 | 0 | 0 | 0 | NB163 | 0 | 0 | 0 | 0 |
| NB170 | 0 | 0 | 1 | 0 | NB164 | 0 | 0 | 0 | 0 | NB165 | 1 | 0 | 0 | 0 | NB167 | 0 | 0 | 0 | 0 |
| NB174 | 0 | 0 | 0 | 0 | NB168 | 1 | 0 | 0 | 0 | NB169 | 0 | 1 | 1 | 1 | NB171 | 0 | 0 | 0 | 0 |
| NB178 | 0 | 0 | 0 | 0 | NB172 | 0 | 0 | 0 | 0 | NB173 | 1 | 0 | 1 | 0 | NB175 | 1 | 1 | 0 | 0 |
| NB182 | 0 | 0 | 0 | 0 | NB176 | 0 | 0 | 0 | 0 | NB177 | 1 | 0 | 1 | 0 | NB179 | 0 | 0 | 1 | 0 |
| NB190 | 0 | 0 | 1 | 1 | NB180 | 1 | 0 | 0 | 0 | NB181 | 0 | 0 | 1 | 0 | NB183 | 0 | 0 | 0 | 0 |
| NB194 | 1 | 1 | 1 | 1 | NB184 | 0 | 0 | 0 | 0 | NB185 | 1 | 0 | 0 | 0 | NB187 | 0 | 0 | 0 | 0 |
| NB198 | 1 | 1 | 1 | 1 | NB186 | 1 | 0 | 1 | 0 | NB189 | 0 | 0 | 1 | 0 | NB191 | 1 | 0 | 1 | 0 |
| NB202 | 0 | 0 | 0 | 0 | NB188 | 1 | 0 | 1 | 1 | NB193 | 1 | 0 | 0 | 0 | NB195 | 1 | 1 | 1 | 1 |
| NB206 | 0 | 0 | 0 | 0 | NB192 | 0 | 0 | 1 | 1 | NB197 | 1 | 1 | 1 | 1 | NB199 | 0 | 0 | 0 | 0 |
| NB210 | 0 | 0 | 0 | 0 | NB196 | 0 | 1 | 0 | 0 | NB201 | 0 | 0 | 0 | 0 | NB203 | 0 | 0 | 0 | 0 |
| NB214 | 1 | 0 | 0 | 0 | NB200 | 0 | 0 | 0 | 0 | NB205 | 0 | 0 | 0 | 0 | NB207 | 1 | 0 | 0 | 0 |
| NB218 | 0 | 0 | 0 | 0 | NB204 | 1 | 0 | 0 | 0 | NB209 | 1 | 0 | 0 | 0 | NB211 | 0 | 0 | 0 | 0 |
| NB226 | 0 | 0 | 0 | 0 | NB208 | 1 | 0 | 0 | 0 | NB213 | 0 | 0 | 0 | 0 | NB215 | 0 | 0 | 0 | 0 |
| NB230 | 0 | 0 | 0 | 0 | NB212 | 0 | 0 | 1 | 0 | NB217 | 0 | 0 | 0 | 0 | NB219 | 0 | 0 | 0 | 0 |
| NB234 | 1 | 0 | 1 | 0 | NB216 | 0 | 0 | 0 | 0 | NB221 | 0 | 0 | 0 | 0 | NB223 | 1 | 0 | 1 | 0 |
| NB238 | 0 | 0 | 0 | 0 | NB220 | 1 | 0 | 0 | 0 | NB225 | 0 | 0 | 0 | 0 | NB227 | 0 | 0 | 1 | 0 |
| NB242 | 1 | 1 | 1 | 1 | NB222 | 1 | 0 | 1 | 0 | NB229 | 1 | 0 | 0 | 0 | NB231 | 0 | 0 | 1 | 0 |
| NB246 | 0 | 1 | 1 | 0 | NB224 | 1 | 0 | 1 | 0 | NB233 | 0 | 0 | 1 | 0 | NB235 | 1 | 0 | 1 | 0 |
| NB250 | 1 | 1 | 1 | 1 | NB228 | 1 | 0 | 0 | 0 | NB237 | 1 | 0 | 0 | 0 | NB239 | 0 | 1 | 0 | 0 |
| NB254 | 0 | 1 | 0 | 0 | NB232 | 1 | 0 | 1 | 0 | NB241 | 0 | 1 | 0 | 0 | NB243 | 0 | 1 | 0 | 0 |
| NB258 | 1 | 1 | 0 | 0 | NB236 | 0 | 1 | 0 | 0 | NB245 | 0 | 1 | 1 | 1 | NB247 | 0 | 1 | 1 | 0 |
| NB262 | 1 | 1 | 1 | 1 | NB240 | 1 | 0 | 1 | 0 | NB249 | 0 | 1 | 0 | 0 | NB251 | 0 | 1 | 0 | 0 |
| NB266 | 1 | 1 | 0 | 0 | NB244 | 0 | 1 | 1 | 1 | NB253 | 1 | 0 | 0 | 0 | NB255 | 1 | 1 | 1 | 1 |
| NB270 | 0 | 1 | 1 | 1 | NB248 | 0 | 1 | 0 | 0 | NB257 | 1 | 1 | 1 | 1 | NB259 | 1 | 1 | 1 | 1 |
| NB274 | 1 | 0 | 0 | 0 | NB252 | 1 | 1 | 1 | 1 | NB261 | 0 | 1 | 0 | 0 | NB263 | 1 | 1 | 1 | 1 |
| NB278 | 0 | 0 | 0 | 0 | NB256 | 1 | 1 | 1 | 1 | NB265 | 0 | 1 | 0 | 0 | NB267 | 1 | 1 | 1 | 1 |
| NB282 | 0 | 0 | 1 | 0 | NB260 | 1 | 1 | 0 | 0 | NB269 | 1 | 1 | 1 | 1 | NB271 | 0 | 1 | 1 | 1 |
| NB286 | 0 | 0 | 0 | 0 | NB264 | 1 | 1 | 1 | 1 | NB273 | 0 | 0 | 1 | 0 | NB275 | 1 | 0 | 0 | 0 |
| NB290 | 0 | 0 | 0 | 0 | NB268 | 0 | 1 | 1 | 1 | NB277 | 1 | 0 | 0 | 0 | NB279 | 0 | 0 | 0 | 0 |
| NB294 | 1 | 0 | 0 | 0 | NB272 | 0 | 1 | 1 | 0 | NB281 | 0 | 0 | 0 | 0 | NB283 | 0 | 0 | 0 | 0 |
| NB298 | 1 | 0 | 0 | 0 | NB276 | 0 | 0 | 0 | 0 | NB285 | 0 | 0 | 0 | 0 | NB287 | 1 | 0 | 1 | 1 |
| NB302 | 1 | 0 | 1 | 0 | NB280 | 1 | 0 | 0 | 0 | NB289 | 1 | 0 | 1 | 1 | NB291 | 1 | 0 | 0 | 0 |
| NB306 | 1 | 0 | 0 | 0 | NB284 | 0 | 0 | 1 | 0 | NB293 | 0 | 0 | 0 | 0 | NB295 | 1 | 0 | 1 | 0 |
| NB310 | 0 | 0 | 0 | 0 | NB288 | 1 | 0 | 0 | 0 | NB297 | 0 | 0 | 1 | 0 | NB299 | 0 | 0 | 0 | 0 |
| NB314 | 0 | 1 | 1 | 1 | NB292 | 0 | 0 | 0 | 0 | NB301 | 0 | 0 | 0 | 0 | NB303 | 0 | 0 | 0 | 0 |
| NB318 | 1 | 0 | 0 | 0 | NB296 | 0 | 0 | 0 | 0 | NB305 | 1 | 0 | 1 | 1 | NB307 | 1 | 0 | 0 | 0 |
| NB322 | 0 | 1 | 1 | 1 | NB300 | 0 | 0 | 1 | 0 | NB309 | 0 | 0 | 1 | 0 | NB311 | 1 | 0 | 1 | 0 |
| NB326 | 1 | 1 | 1 | 1 | NB304 | 0 | 0 | 0 | 0 | NB313 | 0 | 1 | 0 | 0 | NB315 | 0 | 0 | 0 | 0 |
| NB330 | 0 | 0 | 0 | 0 | NB308 | 1 | 0 | 0 | 0 | NB317 | 1 | 0 | 0 | 0 | NB319 | 0 | 1 | 1 | 1 |
| NB334 | 1 | 0 | 0 | 0 | NB312 | 0 | 0 | 0 | 0 | NB321 | 0 | 1 | 1 | 1 | NB323 | 0 | 0 | 1 | 1 |
| NB338 | 1 | 1 | 0 | 0 | NB316 | 0 | 1 | 1 | 1 | NB325 | 0 | 0 | 0 | 0 | NB327 | 0 | 1 | 1 | 1 |
| NB342 | 1 | 0 | 0 | 0 | NB320 | 0 | 1 | 1 | 1 | NB329 | 0 | 0 | 0 | 0 | NB331 | 0 | 1 | 1 | 1 |
| NB346 | 1 | 0 | 0 | 0 | NB324 | 0 | 0 | 0 | 0 | NB333 | 0 | 1 | 1 | 1 | NB335 | 1 | 1 | 1 | 1 |
| NB350 | 1 | 0 | 0 | 0 | NB328 | 1 | 1 | 1 | 1 | NB337 | 0 | 1 | 1 | 1 | NB339 | 1 | 1 | 1 | 1 |
| NB354 | 1 | 0 | 1 | 1 | NB332 | 0 | 1 | 0 | 0 | NB341 | 0 | 1 | 1 | 0 | NB343 | 1 | 0 | 0 | 0 |
| NB358 | 1 | 0 | 1 | 0 | NB336 | 0 | 1 | 1 | 0 | NB345 | 0 | 0 | 0 | 0 | NB347 | 1 | 0 | 0 | 0 |
| NB362 | 0 | 1 | 1 | 1 | NB340 | 1 | 0 | 0 | 0 | NB349 | 1 | 0 | 0 | 0 | NB351 | 0 | 1 | 0 | 0 |
| NB366 | 0 | 0 | 0 | 0 | NB344 | 0 | 0 | 0 | 0 | NB353 | 0 | 0 | 1 | 1 | NB355 | 0 | 0 | 0 | 0 |
| NB370 | 0 | 0 | 0 | 0 | NB348 | 0 | 0 | 0 | 0 | NB357 | 1 | 0 | 1 | 0 | NB359 | 0 | 1 | 1 | 1 |
| NB374 | 0 | 1 | 1 | 0 | NB352 | 0 | 1 | 0 | 0 | NB361 | 0 | 0 | 1 | 0 | NB363 | 0 | 0 | 0 | 0 |
| NB378 | 1 | 1 | 1 | 1 | NB356 | 0 | 0 | 0 | 0 | NB365 | 0 | 0 | 0 | 0 | NB367 | 1 | 0 | 0 | 0 |
| NB382 | 1 | 0 | 0 | 0 | NB360 | 0 | 1 | 1 | 1 | NB369 | 0 | 1 | 1 | 1 | NB371 | 1 | 1 | 1 | 1 |
| NB386 | 1 | 1 | 1 | 1 | NB364 | 0 | 0 | 0 | 0 | NB373 | 1 | 1 | 1 | 1 | NB375 | 0 | 1 | 1 | 1 |
| NB390 | 1 | 1 | 1 | 1 | NB368 | 1 | 0 | 0 | 0 | NB377 | 0 | 1 | 1 | 1 | NB379 | 0 | 1 | 0 | 0 |
| NB394 | 0 | 1 | 1 | 0 | NB372 | 0 | 1 | 1 | 0 | NB381 | 1 | 1 | 1 | 1 | NB383 | 0 | 1 | 1 | 1 |
| NB398 | 0 | 0 | 1 | 0 | NB376 | 0 | 1 | 1 | 1 | NB385 | 1 | 1 | 1 | 1 | NB387 | 0 | 0 | 0 | 0 |
| NB402 | 1 | 1 | 1 | 0 | NB380 | 1 | 0 | 0 | 0 | NB389 | 0 | 1 | 0 | 0 | NB391 | 0 | 1 | 0 | 0 |
| NB406 | 1 | 1 | 1 | 1 | NB384 | 1 | 1 | 1 | 1 | NB393 | 0 | 1 | 1 | 1 | NB395 | 1 | 0 | 0 | 0 |
| NB410 | 1 | 1 | 0 | 0 | NB388 | 0 | 1 | 1 | 1 | NB397 | 0 | 0 | 0 | 0 | NB399 | 0 | 1 | 0 | 0 |
| NB414 | 0 | 1 | 1 | 1 | NB392 | 0 | 1 | 1 | 1 | NB401 | 0 | 1 | 1 | 0 | NB403 | 0 | 1 | 1 | 1 |
| NB418 | 0 | 1 | 0 | 0 | NB396 | 1 | 1 | 1 | 1 | NB405 | 0 | 1 | 0 | 0 | NB407 | 0 | 1 | 1 | 1 |
| NB422 | 0 | 1 | 1 | 0 | NB400 | 0 | 1 | 1 | 0 | NB409 | 0 | 1 | 0 | 0 | NB411 | 1 | 1 | 1 | 0 |
| NB426 | 0 | 1 | 0 | 0 | NB404 | 1 | 1 | 1 | 1 | NB413 | 1 | 1 | 0 | 0 | NB419 | 1 | 0 | 0 | 0 |
| NB428 | 0 | 1 | 0 | 0 | NB408 | 0 | 1 | 1 | 1 | NB415 | 1 | 1 | 1 | 0 | NB423 | 0 | 0 | 1 | 0 |
| NB430 | 0 | 1 | 1 | 0 | NB412 | 0 | 1 | 1 | 0 | NB417 | 0 | 0 | 0 | 0 | NB427 | 0 | 0 | 0 | 0 |
| NB434 | 0 | 0 | 0 | 0 | NB416 | 0 | 0 | 1 | 0 | NB421 | 0 | 0 | 0 | 0 | NB431 | 1 | 1 | 0 | 0 |
| NB436 | 0 | 1 | 0 | 0 | NB420 | 1 | 0 | 0 | 0 | NB425 | 0 | 0 | 0 | 0 | NB435 | 1 | 0 | 0 | 0 |
| NB438 | 0 | 1 | 1 | 1 | NB424 | 1 | 1 | 0 | 0 | NB429 | 1 | 0 | 0 | 0 | NB439 | 0 | 1 | 1 | 1 |
| NB442 | 1 | 1 | 1 | 1 | NB432 | 1 | 1 | 0 | 0 | NB433 | 1 | 0 | 0 | 0 | NB441 | 0 | 0 | 0 | 0 |
| NB446 | 0 | 1 | 1 | 1 | NB440 | 0 | 1 | 1 | 1 | NB437 | 1 | 1 | 1 | 1 | NB443 | 1 | 1 | 1 | 1 |
| NB450 | 0 | 0 | 0 | 0 | NB444 | 0 | 1 | 1 | 1 | NB445 | 0 | 1 | 1 | 1 | NB447 | 0 | 1 | 1 | 1 |
| NB454 | 0 | 1 | 1 | 1 | NB448 | 1 | 1 | 1 | 0 | NB449 | 1 | 1 | 0 | 0 | NB451 | 0 | 0 | 0 | 0 |
| NB458 | 1 | 1 | 1 | 1 | NB452 | 1 | 0 | 0 | 0 | NB453 | 0 | 0 | 0 | 0 | NB455 | 0 | 0 | 0 | 0 |
| NB462 | 1 | 0 | 0 | 0 | NB456 | 0 | 1 | 1 | 1 | NB457 | 0 | 1 | 1 | 1 | NB459 | 0 | 1 | 0 | 0 |
| NB466 | 1 | 1 | 1 | 0 | NB460 | 1 | 1 | 1 | 0 | NB461 | 0 | 1 | 0 | 0 | NB463 | 1 | 0 | 1 | 0 |
| NB470 | 0 | 0 | 0 | 0 | NB464 | 0 | 1 | 1 | 1 | NB465 | 1 | 1 | 0 | 0 | NB467 | 0 | 1 | 0 | 0 |
| NB474 | 1 | 0 | 1 | 0 | NB468 | 0 | 0 | 1 | 0 | NB469 | 0 | 1 | 1 | 0 | NB471 | 1 | 0 | 0 | 0 |
| NB478 | 0 | 1 | 0 | 0 | NB472 | 1 | 0 | 0 | 0 | NB473 | 1 | 0 | 0 | 0 | NB475 | 0 | 1 | 1 | 1 |
| NB480 | 0 | 1 | 0 | 0 | NB476 | 1 | 0 | 0 | 0 | NB477 | 0 | 0 | 0 | 0 | NB479 | 1 | 0 | 0 | 0 |
| NB482 | 0 | 0 | 0 | 0 | NB484 | 1 | 1 | 0 | 0 | NB481 | 0 | 1 | 0 | 0 | NB483 | 1 | 0 | 1 | 0 |
| NB486 | 0 | 1 | 1 | 0 | NB488 | 1 | 0 | 0 | 0 | NB485 | 1 | 1 | 0 | 0 | NB487 | 0 | 1 | 0 | 0 |
| NB490 | 1 | 1 | 1 | 0 | NB492 | 0 | 1 | 1 | 1 | NB489 | 1 | 1 | 0 | 0 | NB491 | 0 | 1 | 1 | 0 |
| NB494 | 0 | 0 | 0 | 0 | NB496 | 0 | 0 | 1 | 0 | NB493 | 0 | 0 | 0 | 0 | NB495 | 0 | 0 | 0 | 0 |
| NB498 | 1 | 0 | 0 | 0 | | | | | | NB497 | 1 | 0 | 0 | 0 | | | | | |
| | **Gender** | **High-risk** | **rogressio** | **death from disease** | | **Gender** | **High-risk** | **rogressio** | **death from disease** | | **Gender** | **High-risk** | **rogressio** | **death from disease** | | **Gender** | **High-risk** | **rogressio** | **death from disease** |
| | **45%** | **37%** | **38%** | **21%** | | **42%** | **35%** | **38%** | **23%** | | **41%** | **34%** | **35%** | **21%** | | **42%** | **35%** | **36%** | **20%** |

| | Maris::Valid | |
|---|---|---|
| Patient_id | progressio | death from disease |
| W001 | 0 | 0 |
| W002 | 0 | 0 |
| W003 | 0 | 0 |
| W004 | 0 | 0 |
| W005 | 0 | 0 |
| W006 | 0 | 0 |
| W007 | 0 | 0 |
| W008 | 0 | 0 |
| W009 | 0 | 0 |
| W010 | 0 | 0 |
| W011 | 0 | 0 |
| W012 | 0 | 0 |
| W013 | 0 | 0 |
| W014 | 0 | 0 |
| W015 | 0 | 0 |
| W016 | 0 | 0 |
| W017 | 0 | 0 |
| W018 | 0 | 0 |
| W019 | 0 | 0 |
| W020 | 0 | 0 |
| W021 | 0 | 0 |
| W022 | 0 | 0 |
| W023 | 0 | 0 |
| W024 | 0 | 0 |
| W025 | 0 | 0 |
| W026 | 0 | 0 |
| W027 | 0 | 0 |
| W028 | 0 | 0 |
| W029 | 0 | 0 |
| W030 | 0 | 0 |
| W031 | 0 | 0 |
| W032 | 0 | 0 |
| W033 | 0 | 0 |
| W034 | 0 | 0 |
| W035 | 0 | 0 |
| W036 | 0 | 0 |
| W037 | 0 | 0 |
| W038 | 0 | 0 |
| W039 | 0 | 0 |
| W040 | 0 | 0 |
| W041 | 0 | 0 |
| W042 | 0 | 0 |
| W043 | 0 | 0 |
| W044 | 0 | 0 |
| W045 | 0 | 0 |
| W046 | 0 | 0 |
| W047 | 0 | 0 |
| W048 | 0 | 0 |
| W049 | 0 | 0 |
| W050 | 0 | 0 |
| W051 | 0 | 0 |
| W052 | 0 | 0 |
| W053 | 0 | 0 |
| W054 | 0 | 0 |
| W055 | 0 | 0 |
| W056 | 0 | 0 |
| W057 | 0 | 0 |
| W058 | 0 | 0 |
| W059 | 0 | 0 |
| W060 | 0 | 0 |
| W061 | 0 | 0 |
| W062 | 1 | 1 |
| W063 | 1 | 1 |
| W064 | 1 | 1 |
| W065 | 1 | 1 |
| W066 | 1 | 1 |
| W067 | 1 | 1 |
| W068 | 1 | 1 |
| W069 | 1 | 1 |
| W070 | 1 | 1 |
| W071 | 1 | 1 |
| W072 | 1 | 1 |
| W073 | 1 | 1 |
| W074 | 1 | 0 |
| W075 | 1 | 1 |
| W076 | 1 | 1 |
| W077 | 1 | 1 |
| W078 | 1 | 0 |
| W079 | 1 | 0 |
| W080 | 1 | 0 |
| W081 | 1 | 1 |
| W082 | 1 | 1 |
| W083 | 1 | 1 |
| W084 | 1 | 1 |
| W085 | 1 | 1 |
| W086 | 1 | 1 |
| W087 | 1 | 1 |
| W088 | 1 | 1 |
| W089 | 1 | 1 |
| W090 | 1 | 1 |
| W091 | 1 | 0 |
| W092 | 1 | 1 |
| | progressio | death from disease |
| | 34% | 28% |

| | Versteeg::Valid | |
|---|---|---|
| Patient_id | progression | ath from disea |
| X001 | 0 | 0 |
| X002 | 0 | 0 |
| X003 | 0 | 0 |
| X004 | 0 | 0 |
| X005 | 0 | 0 |
| X006 | 0 | 0 |
| X007 | 0 | 0 |
| X008 | 0 | 0 |
| X009 | 0 | 0 |
| X010 | 0 | 0 |
| X011 | 0 | 0 |
| X012 | 0 | 0 |
| X013 | 0 | 0 |
| X014 | 0 | 0 |
| X015 | 0 | 0 |
| X016 | 0 | 0 |
| X017 | 0 | 0 |
| X018 | 0 | 0 |
| X019 | 0 | 0 |
| X020 | 0 | 0 |
| X021 | 0 | 0 |
| X022 | 0 | 0 |
| X023 | 0 | 0 |
| X024 | 0 | 0 |
| X025 | 0 | 0 |
| X026 | 0 | 0 |
| X027 | 0 | 0 |
| X028 | 0 | 0 |
| X029 | 0 | 0 |
| X030 | 0 | 0 |
| X031 | 0 | 0 |
| X032 | 0 | 0 |
| X033 | 0 | 0 |
| X034 | 0 | 0 |
| X035 | 0 | 0 |
| X036 | 0 | 0 |
| X037 | 0 | 0 |
| X038 | 0 | 0 |
| X039 | 0 | 0 |
| X040 | 0 | 0 |
| X041 | 0 | 0 |
| X042 | 0 | 0 |
| X043 | 0 | 0 |
| X044 | 0 | 0 |
| X045 | 0 | 0 |
| X046 | 0 | 0 |
| X047 | 0 | 0 |
| X048 | 0 | 0 |
| X049 | 0 | 0 |
| X050 | 0 | 0 |
| X051 | 0 | 0 |
| X052 | 0 | 0 |
| X053 | 0 | 0 |
| X054 | 1 | 1 |
| X055 | 1 | 1 |
| X056 | 1 | 1 |
| X057 | 1 | 1 |
| X058 | 1 | 0 |
| X059 | 1 | 1 |
| X060 | 1 | 0 |
| X061 | 1 | 1 |
| X062 | 1 | 1 |
| X063 | 1 | 1 |
| X064 | 1 | 1 |
| X065 | 1 | 1 |
| X066 | 1 | 1 |
| X067 | 1 | 0 |
| X068 | 1 | 1 |
| X069 | 1 | 1 |
| X070 | 1 | 1 |
| X071 | 1 | 1 |
| X072 | 1 | 1 |
| X073 | 1 | 0 |
| X074 | 1 | 0 |
| X075 | 1 | 1 |
| X076 | 1 | 1 |
| X077 | 1 | 1 |
| X078 | 1 | 1 |
| X079 | 1 | 1 |
| X080 | 1 | 1 |
| X081 | 1 | 1 |
| X082 | 1 | 1 |
| X083 | 1 | 1 |
| X084 | 1 | 1 |
| X085 | 1 | 1 |
| X086 | 1 | 1 |
| X087 | 1 | 1 |
| X088 | 1 | 1 |
| | progression | ath from disea |
| | 40% | 34% |
